# Megaviruses contain various genes encoding for eukaryotic vesicle trafficking factors

**DOI:** 10.1101/2022.01.28.478187

**Authors:** Emilie Neveu, Dany Khalifeh, Dirk Fasshauer

**Affiliations:** Department of Computational Biology, University of Lausanne, Génopode, 1015 Lausanne, Switzerland

**Keywords:** membrane trafficking, SNARE Proteins, Ras Protein, N-ethylmaleimide-sensitive, Factor, Munc18 Proteins, *Mimivirus*, *Legionella*, Giant Viruses, Eukaryota

## Abstract

Many intracellular pathogens, such as bacteria and large viruses, enter eukaryotic cells via phagocytosis, then replicate and proliferate inside the host. To avoid degradation in the phagosomes, they have developed strategies to modify vesicle trafficking. Although several strategies of bacteria have been characterized, it is not clear whether viruses also interfere with the vesicle trafficking of the host. Recently, we came across SNARE proteins encoded in the genomes of several bacteria of the order Legionellales. These pathogenic bacteria may use SNAREs to interfere with vesicle trafficking, since SNARE proteins are the core machinery for vesicle fusion during transport. They assemble into membrane-bridging SNARE complexes that bring membranes together. We now have also discovered SNARE proteins in the genomes of diverse giant viruses. Our biochemical experiments showed that these proteins are able to form SNARE complexes. We also found other key trafficking factors that work together with SNAREs such as NSF, SM, and Rab proteins encoded in the genomes of giant viruses, suggesting that viruses can make use of a large genetic repertoire of trafficking factors. Most giant viruses possess different collections, suggesting that these factors entered the viral genome multiple times. In the future, the molecular role of these factors during viral infection need to be studied.

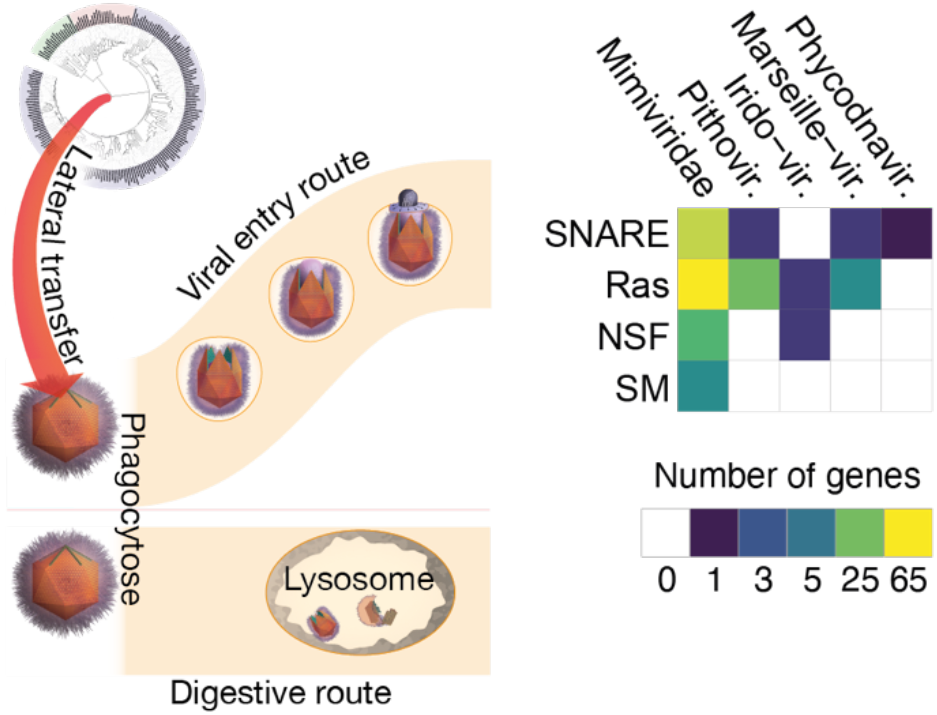

**Synopsis:** Giant viruses enter their eukaryotic host cells by phagocytosis. For reproduction, they hijack the host cell’s membranes by an unknown mechanism. Here, we found that giant viruses express several core factors of the eukaryotic vesicle fusion machinery, including SNARE, Rab, SM proteins, and the disassembly protein NSF. Very probably, these genes were transferred from different eukaryotic hosts to different viruses. Whether the viruses deploy these factors for interfering with the vesicle trafficking of the host cell needs to be investigated.

## Introduction

Eukaryotic cells contain various spatially and functionally separated membrane-bound compartments, each harbouring distinctive sets of lipids and proteins. These macromolecules are newly synthesized in the endoplasmic reticulum (ER); distributed to other cellular compartments such as the Golgi apparatus, lysosomes, and endosomes; and released at the plasma membrane through membrane trafficking. This mechanism also enables the cells to take up material from the extracellular space through the endocytic pathway.

In each trafficking pathway, cargo-loaded vesicles bud from the source membrane and then fuse with a target compartment. Key players in the fusion of a transport vesicle with its acceptor membrane, are the so-called SNARE (soluble N-ethylmaleimide-sensitive factor attachment protein receptor) proteins that fuse membranes through their force-generating assembly mechanism. SNARE complex formation between two membranes is orchestrated by other proteins that belong to conserved families such as Rab small GTPases, Sec1/ Munc18-like (SM) proteins, and tethering proteins. After the membranes have merged, the ATPase N-ethylmaleimide-sensitive factor (NSF) is required to break apart the assembled SNARE complexes[1–5].

These trafficking factors are highly conserved among eukaryotes but are thought to be absent in prokaryotes. To our surprise, however, we recently came across genes encoding for SNARE proteins in γ-proteobacteria of the order Legionellales, including in the well-studied Gram-negative pathogen *L. pneumophila* [6]. These γ-proteobacteria live and replicate inside eukaryotic cells. They enter their eukaryotic host by phagocytosis, an endocytic process that allows diverse eukaryotes to take up large particles via large, actin-driven membrane protrusions. Normally, the phagocytosed material is enzymatically degraded in the vesicles of the endosome/lysosome pathway. In order to escape this fate, *L. pneumophila* injects effector proteins into the host’s cytosol that modulate the activity of trafficking factors such as SNARE and Rab proteins. This prevents the fusion of the phagosome with endosomes and instead promotes the fusion of ER-derived vesicles with the phagosome to produce an ER-like vacuole in which the bacteria can proliferate [7–11].

In 2003, a microorganism visible by light microscopy, and therefore first mistaken for a bacterium living inside the amoeba *Acanthamoeba polyphaga*, was identified as the largest virus discovered at that time [12]. Its virion has a diameter of about 0.7 μM and contains a double-stranded DNA (dsDNA) genome of 1.2 Mb that putatively encodes for more than 900 proteins. This is in stark contrast to most known viruses, which are much smaller and have more streamlined genomes [13]This, which had virus with unprecedented genetic complexity, was named *Acanthamoeba polyphaga mimivirus*, for microbe-mimicking virus, as it enters the host cell like a bacterium via phagocytosis because of its sheer size [14–17]. Note that many smaller, classical enveloped viruses such as influenza virus or SARS-CoV-2 [18] and bacterial toxins such as clostridial neurotoxins [19] enter eukaryotic cells by endocytosis. These viruses and toxins have dedicated docking proteins that bind specifically to cellular surface receptors. Often, the acidic environment of the endosomal vesicles triggers conformational rearrangements of the docking complex, releasing the viral genome [20]or the toxin[21] into the cytosol of the host cell.

The genome of *Mimivirus* is encapsulated by a protein core that is surrounded by a membrane, which is bound by a capsid that is covered by long fibrils. Once phagocytosed, the virus is rapidly delivered to acidified phagosomes, where a unique capsid vertex, referred to as a “stargate” structure, that protrudes from the virion surface opens, exposing the inner lipid membrane of the virus [22]. Fusion of the inner viral membrane with the limiting membrane of the phagosome releases the viral genome into the cytosol [22]. It is unclear yet whether the recently discovered low pH-dependence of *Mimivirus* entry [23] resemble that of the fusion mechanism of classical enveloped viruses. Similar mechanisms have been described for other giant viruses [24]. Subsequent viral replication takes place in elaborate cytoplasmic “viral factories”, where spatial and temporal coordination of virion assembly occurs.

During the last few decades, many other giant viruses have been discovered, revealing that they are common in nature. Giant viruses are grouped together in the recently established, expansive phylum *Nucleocytoviricota*, also known as nucleocytoplasmic large DNA viruses [25][26]. They are subdivided into several families that exhibit a broad host range from single cell amoebae (e.g. *Mimivirus*) to humans (e.g. *Poxviridae*). The phylum name stems from the fact that these viruses replicate in viral factories in the cytoplasm of the host cell, although several substantial morphological differences between virus families have been described [16]. Analogous to intracellular bacteria, the establishment of the elaborate viral factories involves the recruitment of host factors and massive rearrangements of cellular membranes.

A crucial step in virus assembly is the formation of inner membranes. Initial work on poxviruses demonstrated that they originate from the host cell’s ER. At the later stages of *Mimivirus* assembly, ER-derived vesicles of ≈ 70 nm are recruited to the periphery of the viral factors, where they fuse into multivesicular bodies that give rise to the inner viral membrane surrounding the viral core [27] Treatment with Brefeldin A, a membrane trafficking inhibitor, has been shown to negatively affect the morphogenesis of virus particles in several different giant viruses [28–31], corroborating the idea that vesicle trafficking in the host cell is important for virus formation. At the endpoint of infection, the newly formed virions fill the host cell and are released into the environment upon its lysis.

Alongside to a core of about 40 viral genes, giant viruses have acquired many genes from different domains of cellular life during their evolution [32]. Their genome encodes for proteins previously considered as signatures of cellular organisms such as DNA methyltransferases, DNA site-specific endonucleases, glycosylating enzymes, and numerous DNA polymerase subunits. In contrast to intracellular bacteria, the viral and cellular factors involved in these virus-induced membrane trafficking and rearrangement events are still largely unknown.

A few years ago, it was reported that the *Mimivirus* genome encodes a Rab GTPase that might play a role in redirection of the secretory pathway, such as the hijacking of ER-derived vesicles during virus production [33]. It is unclear, however, whether other eukaryotic trafficking factors play a role as well during viral infection and production. When we established a classification of SNARE proteins, we came across two different SNARE proteins encoded in the genome of viruses [34], an R-SNARE encoded by *Coccolithovirus* [35] and a Qbc-SNARE in the genome of *Mimivirus* [36]. In 2006, the detection of two viral sequences in ≈ 3600 eukaryotic SNARE protein sequences was only a side note [34]. However, in light of the discovery of multiple SNARE genes in the genomes of intracellular bacteria [6], we have now extended our search for SNAREs and other trafficking factors in giant viruses.

## Results

In order to search for viral SNARE proteins, we scanned the NCBI protein databases for viruses using hidden Markov model (HMM) profiles trained previously to classify eukaryotic SNARE proteins [34]. As in our previous study [6], we implemented a 1E^-4^ expectation value cutoff to minimize false positive results. We found about 80 true SNARE sequences encoded in the genomes of various different giant viruses. Based on their sequence profiles, they were assigned with ease to the four basic types, namely Qa-, Qb-, Qc-, and R-SNAREs. These basic types reflect their position in the heterologous four-helix bundle that is formed during vesicle fusion in eukaryotic cells. In addition, we found several viruses, including *Mimivirus*, that encodes for SNAP-25-like (25-kDa synaptosome-associated protein) proteins containing two SNARE motifs, a so-called Qbc-SNAREs. Qbc-SNAREs are thought to work in endosomal trafficking and secretion. Although some giant virus have only one SNARE protein, some viruse genomes encode for several SNAREs. The highest number of SNARE genes – eight - was found in *Yasminevirus*. Giant viruses encoding SNARE genes are given in Figure 1.

**Figure 1.**
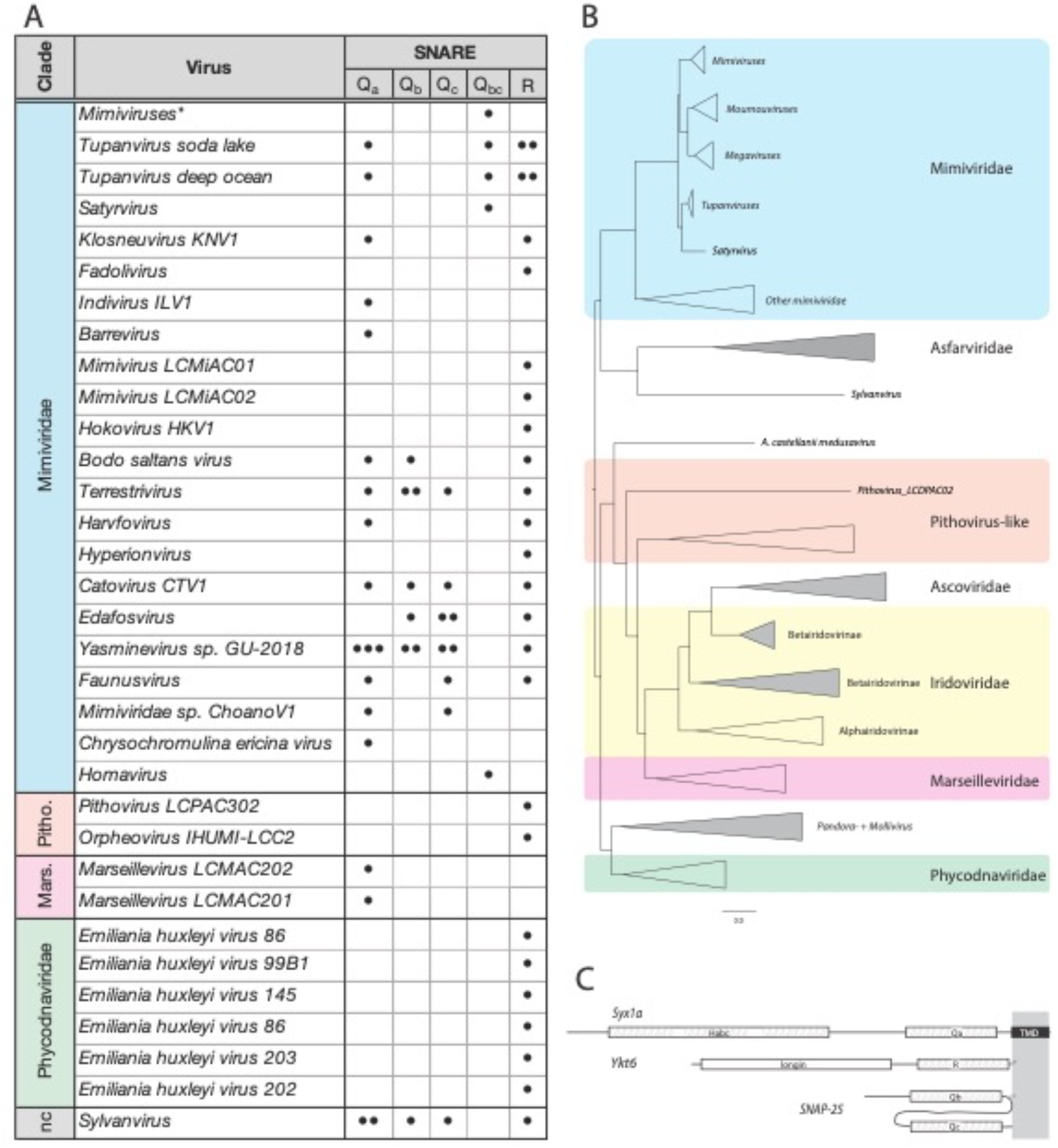
Schematic overview of giant virus SNARE proteins. A) Overview of the distribution and types of SNARE proteins (black circles) in giant viruses. The corresponding sequence IDs are given in Table S2. B) Schematic tree of giant viruses based on an alignment of DNA polymerase B, a protein shared among giant viruses [62]. The tree was calculated using IQ-TREE. Scale bar indicates the number of substitutions per site. The tree was visualized using FigTree v1.4.4 (https://github.com/rambaut/figtree/releases). C) The domain architecture of exemplary SNARE proteins (Syx1a, Ykt6, and SNAP-25 from *Homo sapiens* are shown). Similar SNARE protein types were found in giant viruses.

To check for close homologies, we clustered viral SNAREs together with previously discovered bacterial SNAREs [6] using the basic local alignment search tool (BLAST) to construct homology groups for further inspection. Although we retrieved the clusters of bacterial SNAREs described previously (see also below), most viral SNAREs were not found in larger clusters. One noteworthy cluster of viral sequences contained closely related R-SNARE sequences of different *Emiliania huxleyi* viruses [33].

In preliminary phylogenetic surveys, viral SNARE genes clustered with different eukaryotic lineages, in most cases, no close relationship between viral and eukaryotic sequences materialized. Exceptions were the R-SNAREs from several *Emiliania huxleyi* viruses, which clustered with endosomal R-SNAREs from haptophytes such as *Emiliania huxleyi* and *Chrysochromulina tobinii*. Another example is the R-SNARE of the *Bodo saltans virus* (ATZ81026.1), which clustered with R-SNARE sequences from kinetoplastids, suggesting that it was acquired by the virus from a host belonging to that eukaryotic lineage. As most viral SNARE protein sequences are often quite divergent, it was not possible to assign them unambiguously to a distinct trafficking step, although many appear to belong to SNARE protein types that work in endosomal trafficking.

In an earlier study, we found a comparable distribution of SNARE types in the SNARE inventory of γ-proteobacteria of the order Legionellales [6]. With the exception of *Berkiella cookevillensis* [41], whose metagenome codes for three different SNARE proteins (an R-, a Qbc-, and a Qa-SNARE) most Legionellales bacteria have only one SNARE protein [6]. We also reported a cluster of homologous R-SNAREs in the genomes of various species in the genus *Legionella* [6]. Another cluster contained Qc-SNAREs, termed LseA[37], from various strains of *L. pneumophila*. Two homologous Qbc-SNAREs were found in the genomes of *L. cherrii* and *L. gormanii*. A Qa-SNARE was found in *L. santicrucis*, revealing a broad distribution of different SNARE protein types within a relatively small group of related bacteria [6].

We scanned the NCBI protein database again for bacteria to update our previous inventory of SNAREs from γ-proteobacteria of the order Legionellales. Ten additional bacterial SNARE sequences were retrieved (Table S1). Most of them are closely related to a cluster of bacterial R-SNAREs described in our previous study [6]. Two of these bacterial R-SNAREs were unusual, as they contain two R-SNARE motifs in a row; however, whether that domain arrangement is functional is unclear, however. The bacterial R-SNAREs possess a C-terminal CAAX motif, suggesting that they are farnesylated after being synthesized in the cytosol of the bacterium and then translocated by the bacterial Dot/Icm Type IV secretion system into the host cell’s cytosol. In fact, as previously reported, most bacterial SNAREs, independent of their type, do not have a transmembrane domain (TMD). This is in contrast to most eukaryotic SNARE proteins, which are anchored in the membrane via a C-terminal single-pass TMD facing the cytosol. A notable exception is the Qbc-SNAREs, which are often attached to the membrane by palmitoylation of a central cysteine, whereas R-SNAREs of type II ( Ykt6) are another one. Few viral SNAREs were found to have a CAAX motif, while the majority of viral SNARE proteins have a TMD. Note that many proteins (≈ 10 - 40%) encoded by giant viruses have TMDs [16]This probably reflects the fact that the viral genes are translated directly in the host cell’s cytosol and can then be inserted via their membrane anchors into the membrane of a host compartment. In contrast to intracellular bacteria, no complex protein translocation system is required for viral factors.

Overall, the broad phyletic distribution of the SNARE proteins of giant viruses suggests that they have been acquired repeatedly by lateral gene transfer of different SNARE genes from their eukaryotic hosts.

### Viral SNARE proteins can assemble into SNARE complexes in vitro

In order to validate the classification of viral SNARE proteins, we tested the ability of some to form a ternary SNARE complex. For this, we recombinantly expressed and purified four representative viral proteins, a Qbc-SNARE, two R-SNARE and one Qa-SNARE. For binding experiments, the viral SNARE proteins were mixed with the neuronal SNARE proteins Syx1 (Qa-), SNAP-25 (Qbc-), and Syb2 (R-SNARE) in combinations that ensured that a Qa, a Qbc, and an R-SNARE were combined each time. The neuronal SNARE complex assembles during synaptic secretion [3]. It is resistant to SDS, a feature widely used for monitoring complex formation. We chose this particular SNARE unit, because its subunits can interact, to some extent, with SNARE proteins working in other trafficking steps [6]. The complex of Syx1, Syb2, and the Qbc-SNARE from *Mimivirus* was SDS-resistant, as demonstrated by the appearance of a protein band with apparent molecular masses corresponding to ternary complexes (Figure 2). The ternary SNARE complexes of the other three viral SNAREs were not SDS-resistant, but formed stable complexes, as shown by by native gel electrophoresis [27, 29], a technique that allows for the separation of the individual proteins from complexes, provided that the latter are of sufficient stability (Figure S1).

**Figure 2.**
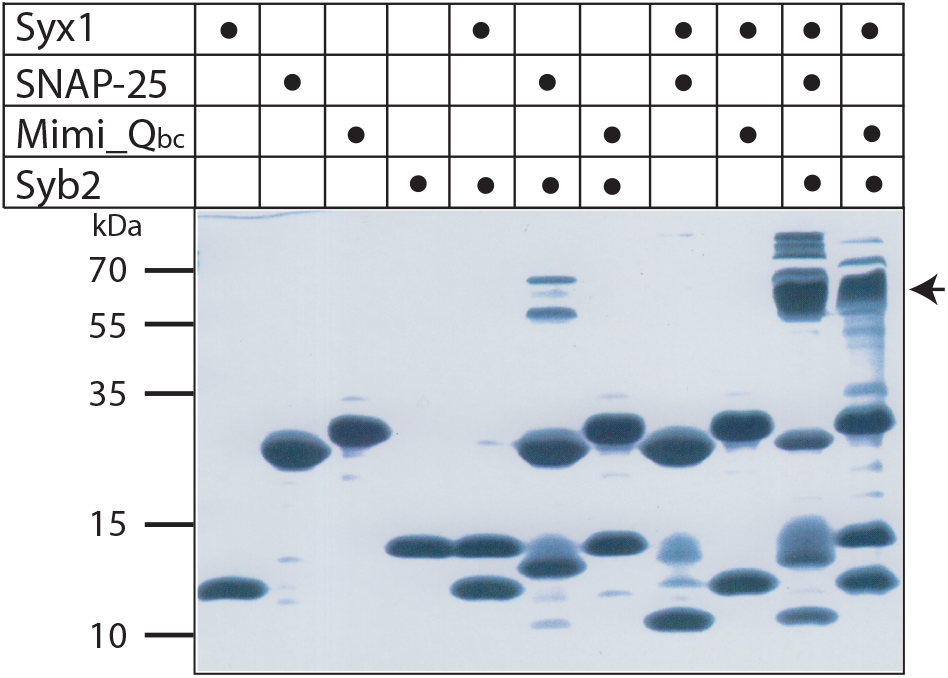
Formation of an SDS-resistant complex between the SNAP-25-like SNARE protein from *Mimivirus* and neuronal SNARE proteins from rat. Approximately equal molar ratios of purified SNARE proteins were mixed, incubated overnight, and subjected to SDS-PAGE without boiling of the sample. After the run, the gel was stained with Coomassie Blue. Neuronal synaptobrevin 2 (Syb) and the SNARE domain of syntaxin 1a (Syx1a) form ternary SDS-resistant complexes (arrow) with neuronal SNAP-25 and with the SNAP-25-like SNARE protein from *Mimivirus*.

### Small GTPases are encoded in the genomes of giant viruses

As outlined above, SNARE proteins form the core of molecular machinery that catalyses the merger of transport vesicles with a target membrane. SNAREs work together with several other conserved proteins such as Rab small GTPases, SM proteins, tethering factors, and the ATPase N-ethylmaleimide-sensitive factor (NSF) that disassembles SNARE complexes after membranes merge (Figure 3). The presence of SNARE proteins in the genome of megaviruses suggests that these viruses may have incorporated other trafficking factors as well.

**Figure 3.**
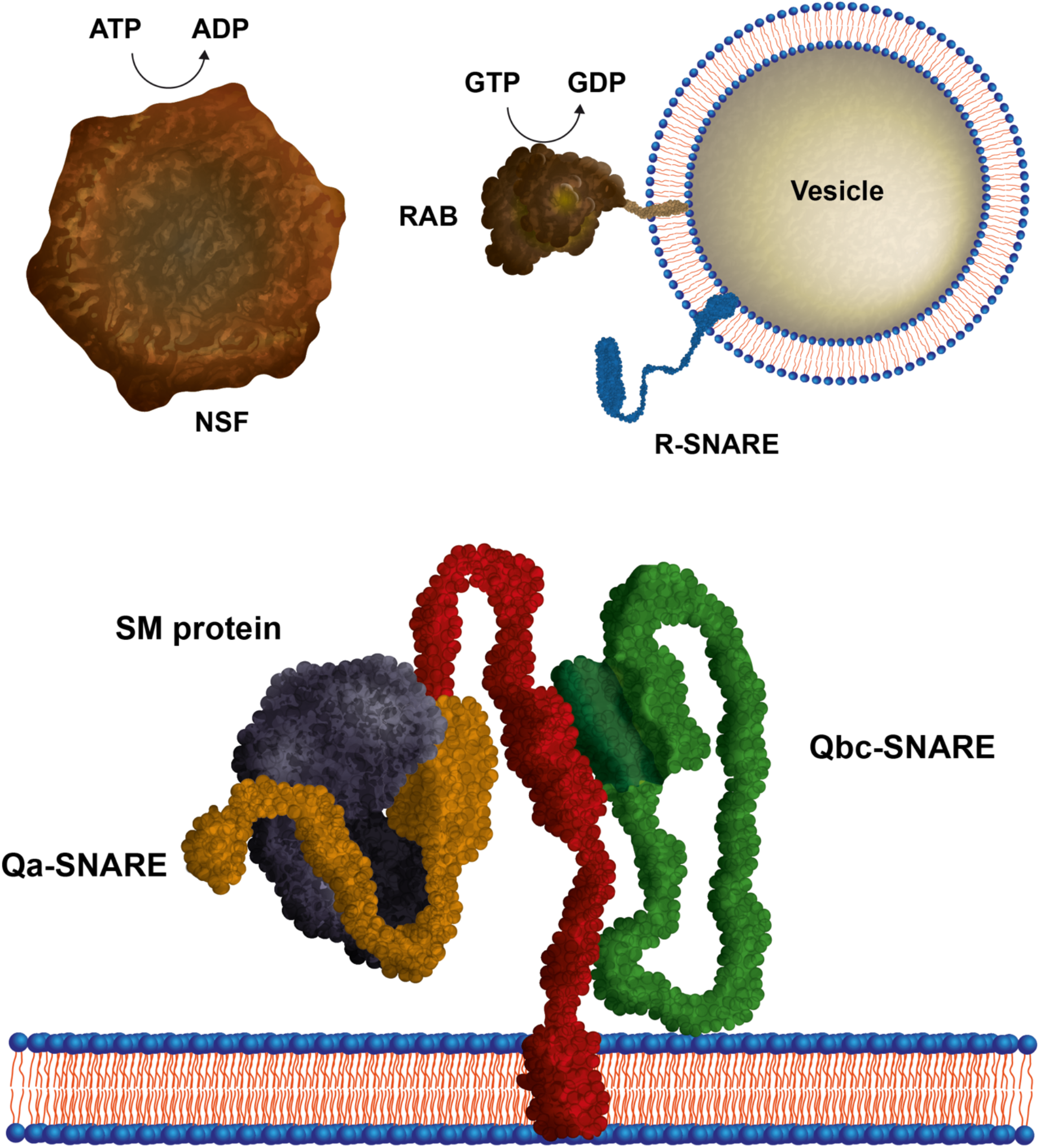
Schematic depiction of key proteins involved in docking and fusion of transport vesicles. The proteins involved in the principal aspects of vesicular trafficking are highly conserved among all eukaryotes, not only among different species but also among different trafficking steps within the cell. At each trafficking step, the vesicle fusion machinery consists of the following: a core of SNARE proteins, which assemble into a tight complex between the membranes, and various other conserved factors, including Sec1/Munc18, Rab, the disassembly ATPase NSF, and other regulatory proteins recruited during different phases of the reaction that orchestrates SNARE activity.

Several years ago, a Rab GTPase homologue was found in the genome of *Mimivirus* and closely related viruses [33]. In a follow-up study, the structure of this *Mimivirus* Rab protein was elucidated. It shows homology to Rab proteins playing a role in endosomal trafficking such as Rab5, Rab22, and Rab21, i.e., belonging to Group II according to our earlier classification of Rab proteins [38]. Rab proteins are thought to specify the identity of vesicles and organelles in order to ensure the specificity of the membrane trafficking steps taken by other recruiting trafficking proteins [39].

In a more recent study, Ran-like GTPases and some other types were found in *Medusavirus* and other viruses [40], suggesting that small GTPases are more widespread in giant viruses. To establish a more complete overview, we next searched next with our HMMs [41] for other Rab-like proteins in viruses. Alongside the Rab protein of *Mimivirus* and related viruses, we uncovered various different Rabs in numerous lineages of megaviruses (Table 1). As well as Rab proteins, we also found other small GTPases of the Ras superfamily, including Ras-, Rho-, and Ran-like sequences. The highest number of small GTPases (11) was found to be encoded in *Terrestrivirus*. We noted that Rab-like GTPases were more often found in the Mimiviridae, whereas most Ran-like GTPases were present in the Pithoviridae.

**Table 1.**
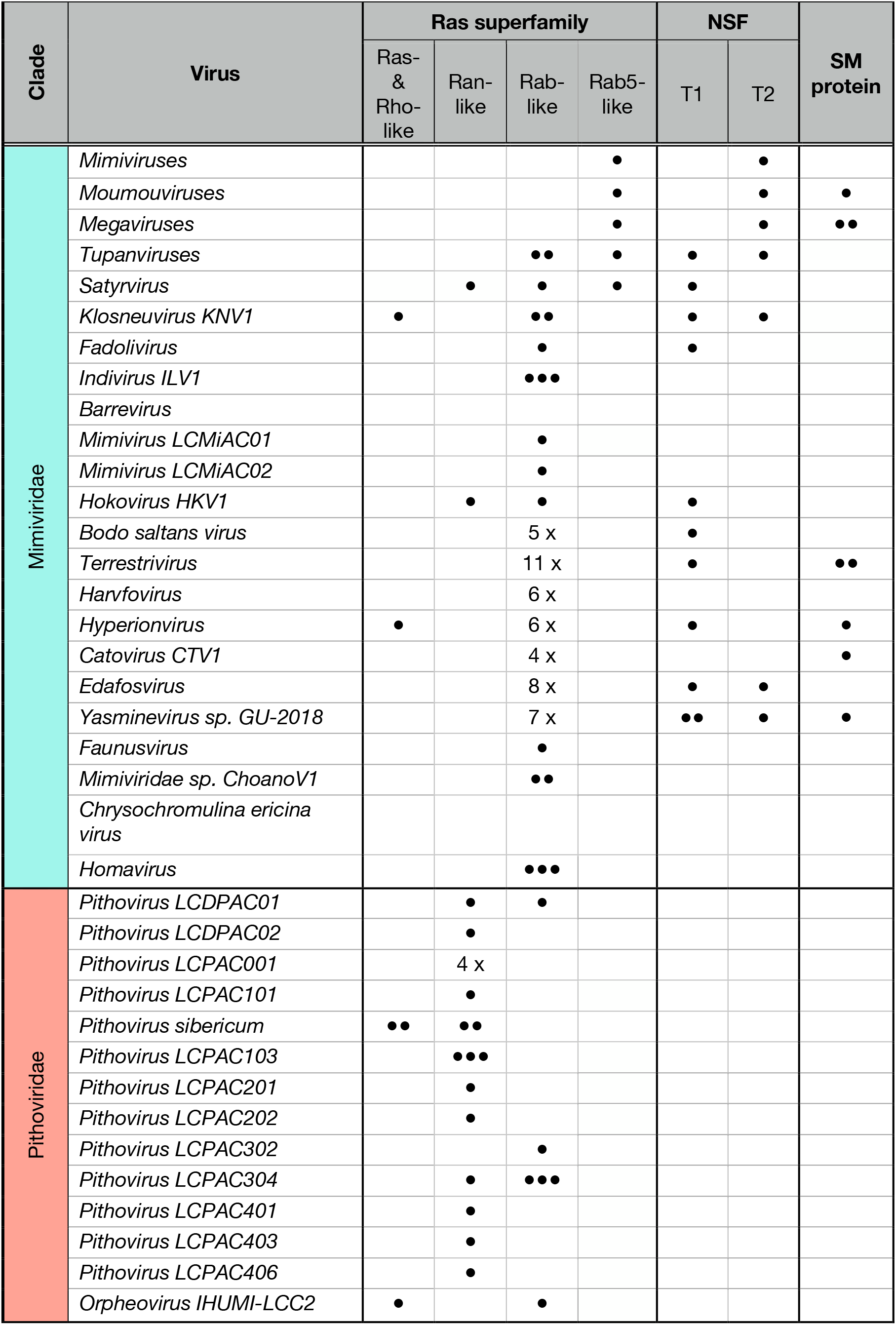
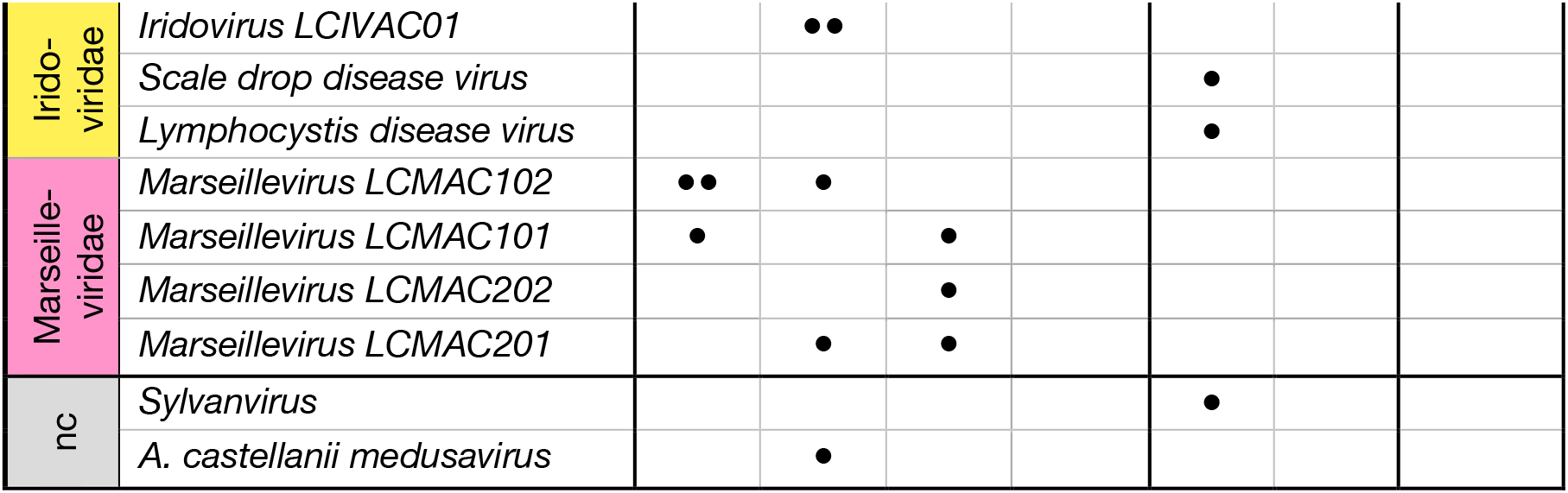
Distribution of other key trafficking factors in the genomes of giant viruses. The presence and number of genes is indicated by black circles or by the number of genes. The corresponding sequence IDs are given in Table S2. nc: not classified.

| Clade | Virus | Ras superfamily |  |  |  | NSF |  | SM protein |
| --- | --- | --- | --- | --- | --- | --- | --- | --- |
|  |  | Ras- & Rho-like | Ran-like | Rab-like | Rab5-like | T1 | T2 |  |
| Mimiviridae | *Mimiviruses* |  |  |  | • |  | • |  |
|  | *Moumouviruses* |  |  |  | • |  | • | • |
|  | *Megaviruses* |  |  |  | • |  | • | •• |
|  | *Tupanviruses* |  |  | •• | • | • | • |  |
|  | *Satyrvirus* |  | • | • | • | • |  |  |
|  | *Klosneuvirus KNV1* | • |  | •• |  | • | • |  |
|  | *Fadolivirus* |  |  | • |  | • |  |  |
|  | *Indivirus ILV1* |  |  | ••• |  |  |  |  |
|  | *Barrevirus* |  |  |  |  |  |  |  |
|  | *Mimivirus LCMiAC01* |  |  | • |  |  |  |  |
|  | *Mimivirus LCMiAC02* |  |  | • |  |  |  |  |
|  | *Hokovirus HKV1* |  | • | • |  | • |  |  |
|  | *Bodo saltans virus* |  |  | 5 x |  | • |  |  |
|  | *Terrestrivirus* |  |  | 11 x |  | • |  | •• |
|  | *Harvfovirus* |  |  | 6 x |  |  |  |  |
|  | *Hyperionvirus* | • |  | 6 x |  | • |  | • |
|  | *Catovirus CTV1* |  |  | 4 x |  |  |  | • |
|  | *Edafosvirus* |  |  | 8 x |  | • | • |  |
|  | *Yasminevirus sp. GU-2018* |  |  | 7 x |  | •• | • | • |
|  | *Faunusvirus* |  |  | • |  |  |  |  |
|  | *Mimiviridae sp. ChoanoV1* |  |  | •• |  |  |  |  |
|  | *Chrysochromulina ericina virus* |  |  |  |  |  |  |  |
|  | *Homavirus* |  |  | ••• |  |  |  |  |
| Pithoviridae | *Pithovirus LCDPAC01* |  | • | • |  |  |  |  |
|  | *Pithovirus LCDPAC02* |  | • |  |  |  |  |  |
|  | *Pithovirus LCPAC001* |  | 4 x |  |  |  |  |  |
|  | *Pithovirus LCPAC101* |  | • |  |  |  |  |  |
|  | *Pithovirus sibericum* | •• | •• |  |  |  |  |  |
|  | *Pithovirus LCPAC103* |  | ••• |  |  |  |  |  |
|  | *Pithovirus LCPAC201* |  | • |  |  |  |  |  |
|  | *Pithovirus LCPAC202* |  | • |  |  |  |  |  |
|  | *Pithovirus LCPAC302* |  |  | • |  |  |  |  |
|  | *Pithovirus LCPAC304* |  | • | ••• |  |  |  |  |
|  | *Pithovirus LCPAC401* |  | • |  |  |  |  |  |
|  | *Pithovirus LCPAC403* |  | • |  |  |  |  |  |
|  | *Pithovirus LCPAC406* |  | • |  |  |  |  |  |
|  | *Orpheovirus IHUMI-LCC2* | • |  | • |  |  |  |  |

|  |  |  |  |  |  |  |
| --- | --- | --- | --- | --- | --- | --- |
| Irido-<br>viridae | *Iridovirus LCIVAC01* |  | •• |  |  |  |
|  | *Scale drop disease virus* |  |  |  |  | • |
|  | *Lymphocystis disease virus* |  |  |  |  | • |
| Marseille-<br>viridae | *Marseillevirus LCMAC102* | •• | • |  |  |  |
|  | *Marseillevirus LCMAC101* | • |  | • |  |  |
|  | *Marseillevirus LCMAC202* |  |  | • |  |  |
|  | *Marseillevirus LCMAC201* |  | • | • |  |  |
| nc | *Sylvanvirus* |  |  |  |  | • |
|  | *A. castellanii medusavirus* |  | • |  |  |  |

Some of the small GTPases of different viruses were found to be related, such as the aforementioned Rab5-like, which is present in many Mimiviridae, and, therefore were placed into a separate column in Table 1. A few other viral small GTPases were found in smaller clusters. For example, a group of three or four Rab proteins of *Harvfovirus* (AYV81889 - AYV81892), *Hyperionvirus* (AYV83656 - AYV83658), and *Edafovirus* (AYV77911 - AYV77914) appear to be related. Interestingly, these Rabs are located next to each other in the viral genomes, suggesting they arose through internal gene duplication events. Note that we also came across a group of trafficking factors located on a stretch in the genome of *Yasminevirus*, where a Rab gene (VBB19039) is located in the vicinity of four SNARE genes (VBB19031, VBB19034, VBB19035, and VBB19046) and one SM protein gene (VBB19033, see below). Most viral Rab proteins, however, are not directly related and even belong to different Rab subtypes [38].

### Other trafficking proteins are encoded in the genomes of giant viruses

When we continued our survey for other components of the vesicle fusion machinery, we came across Sec1/Munc18 (SM) proteins in the genome of several Mimiviridae. SM proteins tightly regulate SNARE complex formation between fusion membranes through clasping a member of the Qa-SNARE subfamily (Figure 3). The viral SM protein sequences are quite divergent and probably belong to either the Vps45 or Vps33 subfamily of SM proteins, which play roles in endosomal trafficking steps. Some megaviruses even encode for two different SM proteins (Table 1).

Breaking apart highly stable SNARE complexes after membrane fusion requires the activity of the ATPase NSF (Figure 3). Using our specific HMMs[42], we found two different types of NSF-like proteins in the genomes of megaviruses (Table 1). One type has the canonical domain architecture found in eukaryotic NSF: an N-terminal domain, which interacts with the SNARE complex, and an adaptor protein, is followed by two ATP-binding domains, termed the D1- and D2-domains. As the catalytic regions of the D-domains of these proteins are conserved, it is conceivable that this viral NSF might function in SNARE complex disassembly, but this would need to be tested biochemically. A survey in BLAST suggested that the viral NSFs of Type 1 may have been transferred repeatedly from different eukaryotic hosts. Note in this respect that the γ-proteobacterium *Legionella santicrucis* also possesses a related NSF gene of Type 1 (WP_058515092.1). Two other putative bacterial NSF were found through a gut metagenome project in *Clostridial bacterium* (MBP3801609.1) and in *Acholeplasmatales bacterium* (MBR4496047.1).

A second type that is more divergent and most probably monophyletic, contains only a D1 domain; it is unclear whether this NSF-like factor is able to bind to and disassemble SNARE complexes. Remarkably, we found a putative homologue of this more divergent NSF-type in the pathogenic bacteria *Waddlia chondrophila* (WP_013181706.1), *Estrella lausannensis* (WP_098039298.1), *Criblamydia sequanensis* (WP_041018683.1), and two other Chlamydiae bacteria (NGX58071.1, NGX58268.1). Whereas the NSF-like protein of *Waddlia* has only one D-domain, the other bacterial proteins appear to have another D-domain *C*-terminally. The region of homology between these Type 2 viral and *Chlamydiae* NSF is not restricted solely to the D1-domain but extends to the N-terminal portion of the sequence as shown in Figure 4.

**Figure 4.**
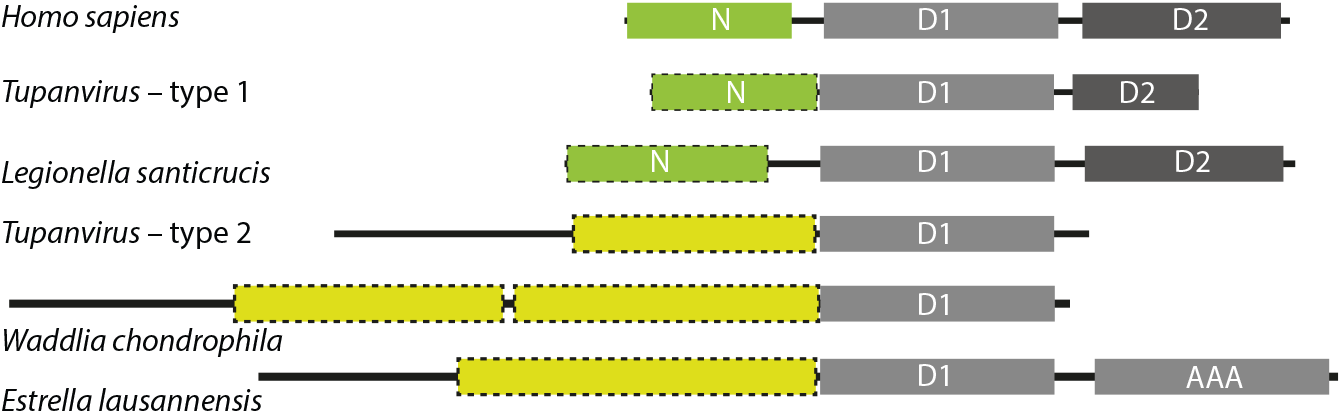
Domain organization of human NSF and different viral and bacterial NSF-like proteins. The tandem D-domains, D1 and D2, are shown in grey. N-domains are shown in green. The conserved N-terminal region of the more divergent type of NSF is highlighted by dashed yellow boxes.

## Discussion

Giant viruses enter their host through phagocytosis [14–17] (Figure 5). After fusion of the viral membrane with the phagosome membrane, the viral genome is released into the cytosol of the host cell and the virus takes over control. Careful morphological studies using electron microscopy have revealed that initial viral replication points gradually merge into a new organelle-like structure, the virus factory, which are often made of membranes of the ER. Mitochondria and the cytoskeleton are often recruited to viral factories [24,43] [16]. Although the morphology and pathways of viral production processes vary somewhat among different giant virus lineages, factors encoded by the virus probably exploit the membrane trafficking routes of their host cells to ensure viral proliferation. That membrane trafficking plays a role during viral replication was also demonstrated by the addition of Brefeldin A, a drug that inhibits vesicle transport from the ER to the Golgi complex [29,30,44]

**Figure 5.**
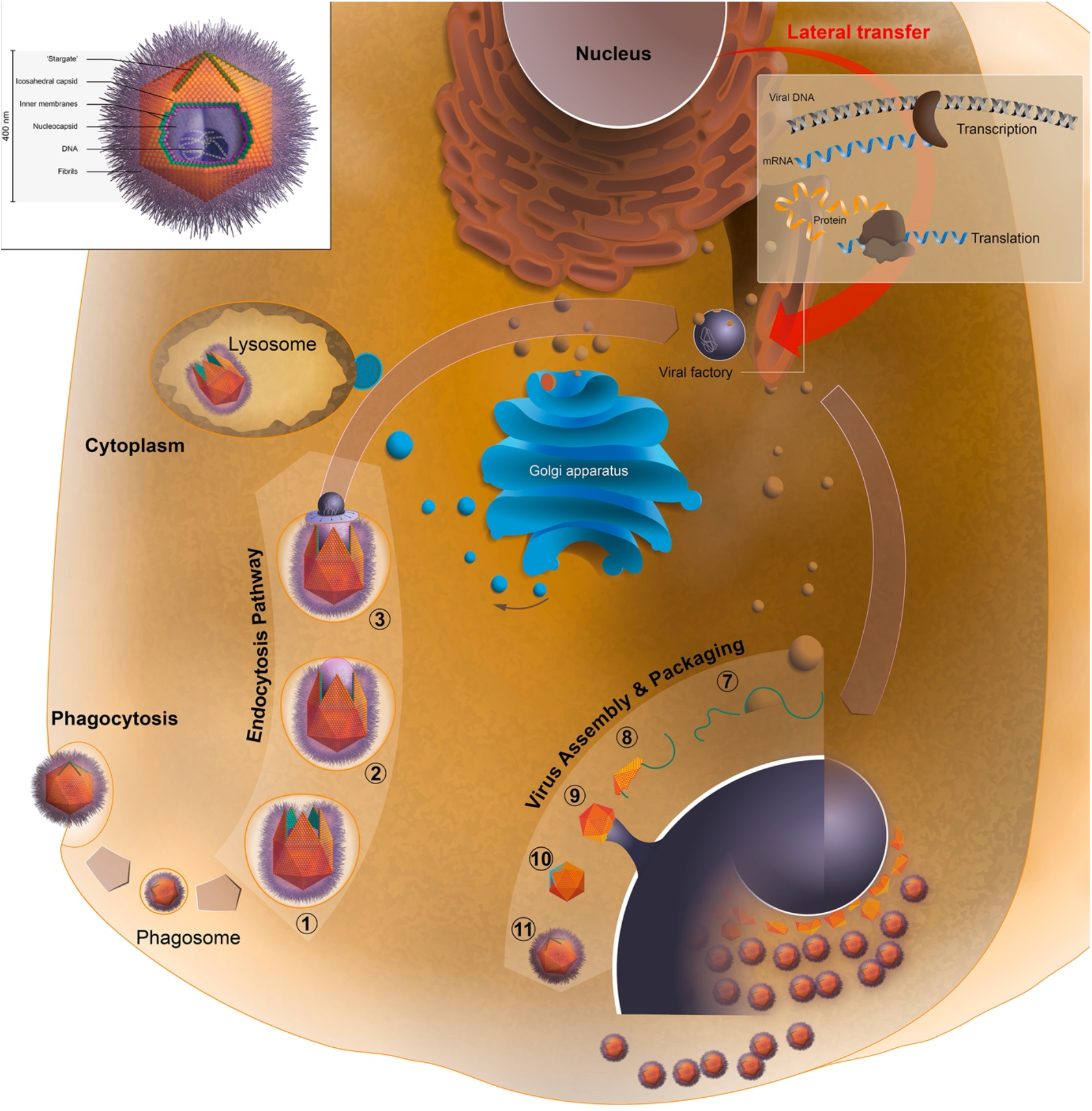
Schematic depiction of the key steps of the entry and replication of giant viruses inside a eukaryotic host cell. A giant virus (the example here is the widely studied *Mimivirus*), is engulfed by phagocytosis. The genome is encapsulated by a protein core that is surrounded by a membrane, which is bound by a capsid covered by long fibrils. In the acidified phagosome, the virus opens and eventually releases its genome through the so-called stargate structure [23] [24]. Upon fusion with the phagosomal membrane, the genome is then released into the host’s cytoplasm [22]. The viral replication, assembly, and packaging occurs in elaborate viral factories that are built from membranes of the host cell. The final viruses are assembled, packed and released from the bursting host cell. Giant viruses contain various genes of eukaryotic origin that may have been incorporated into their genomes by lateral gene transfer during infections.

However, which factors contribute and what strategies are used by the viruses are not clear yet. By contrast, various strategies used by intracellular pathogenic bacteria, which also enter eukaryotic cells by phagocytosis, to redirect the vesicle trafficking of their host cells have been described in much detail [45][46][47][10]. Intracellular bacteria inject proteins produced within the bacterium through their secretion systems into the cytosol of the host cell. Some bacterial effector proteins were acquired by lateral gene transfer and still resemble homologous host proteins, including proteins that interfere with the GTP/GDP cycle of Rab proteins, key proteins for the correct targeting of transport vesicles[48,49]. Recently, *Legionella* were reported to encode for Rab-like proteins as well [50]. In our own studies, we came across several types of SNARE proteins encoded in the genome of γ-proteobacteria of the order Legionellales[6][37], suggesting that these bacteria use these proteins as well to interfere with vesicle trafficking of the host cell.

SNARE proteins are the core machinery for catalysing the fusion of a transport vesicle with the target membrane [1–5]. When we extended our search to other cellular pathogens, i.e. viruses, we discovered that SNARE proteins are also encoded in the genome of giant viruses. The large genomes of giant viruses contain a large number of various genes of eukaryotic origin[17]. These viruses might thus use these proteins as well to interfere with vesicle trafficking. Similar to γ-proteobacterial SNARE proteins [6], the viral SNARE proteins belonged to different SNARE protein types that work in different trafficking steps in eukaryotes. We found viral SNARE proteins in different lineages of giant viruses, and phylogenetic analysis revealed that only a few viral SNARE proteins in closely related viruses were directly related. It was not possible to precisely determine the eukaryotic organism from which the viruses took the SNARE genes, but preliminary BLAST searches indicate that they came from different eukaryotic lineages.

The genomes of giant viruses were reported earlier to code for small GTPases of the Ras superfamily, particularly Rab proteins [33] [38]Together with other small GTP binding protein families such as Ras, Rho, Ran, and Arf, they form the Ras superfamily. Each of these families plays important roles in eukaryotic cells [51]. These GTPases serve as molecular switches and, depending on their status (GTP- or GDP-bound) other effector proteins can interact with them. That a virus can encode for a Ras (“Rat sarcoma”) protein is not new, since the first members of the protein family were identified through studies of cancer-causing viruses, the Harvey and Kirsten sarcoma viruses [52,53].When we scanned virus genomes for Ras sequences, we found the original Ras sequences from these retroviruses and also discovered multiple other small GTPases in the genomes of giant viruses. As noted for the viral SNARE proteins, the small GTP binding proteins are of different eukaryotic types and cannot stem from only one single lateral gene transfer event. Most of the small GTPases are Rab proteins, but several Ran and a few Ras/Rho-like factors are among them as well. Viral Rab proteins belong to different eukaryotic subtypes [41] suggesting that they are distributed to different transport vesicles in the host cell upon virus infection.

Ran proteins, which are essential for the translocation of RNA and proteins through the nuclear pore complex, are mostly found in Pithoviridae, whereas Rab proteins are more common in Mimiviridae. Their different distributions in viral genomes could indicate whether or not the nucleus plays an important role in viral replication. It might also reflect differences in how and from which organelle the virus recruits membranes for its envelope in the host cell.

When we broadened our search, we came across NSF and SM-proteins, other factors of the core vesicle fusion machinery, in the genomes of giant viruses. One viral NSF type closely resembles eukaryotic NSF, the key protein that disassembles SNARE complexes. This canonical NSF probably stems from lateral gene transfer from different eukaryotic hosts. A second, more divergent type of NSF has an altered domain arrangement. Related sequences were found in intracellular bacteria of the phylum Chlamydiae, suggesting that genetic material was exchanged not only between the eukaryotic host and the pathogens but also between different pathogens infecting eukaryotic cells. The amount of gene exchange between different domains of life and its contribution to evolutionary innovation is under investigation [54][55][56]

There seems to be no clear pattern in the distribution of eukaryotic trafficking factors in the genome of giant viruses, but it is interesting to note that some viruses, such as *Yasminevirus* encode for a large repertoire of trafficking factors, whereas the best-studied virus, *Mimivirus* encodes for only few, a Qbc-SNARE, a Rab protein, and an NSF-like factor. However, when and how do the viruses make use of their different sets of trafficking factors? For most factors, we can only assume that they are functional because of their sequences patterns. For example, we noted that the active sites in the GTPases and in NSF seem to be intact. Most viral SNAREs have a transmembrane region and are probably inserted into the membranes of host cell compartments upon translation. Our biochemical data suggest that they can form tight SNARE complexes. The question remains whether they act, during viral entry or if they are part of the viral subversion strategy. Our survey does not answer this question, but a few other studies have provided first insights into the time of activity of viral proteins. The SNARE protein of *Mimivirus* (L657) is only expressed after several hours (3–9h) of viral infection [57], similar to the R-SNARE protein of *Coccolithovirus* [35]. The *Mimivirus* Rab protein (R214) is expressed after 6–9h, but at a much higher level than the SNARE protein [57]. Note that, because of the high divergence of the viral sequences, our HMMs do not allow us to predict precisely which trafficking route in the host cell might be affected by the viral trafficking factors.

The Rab protein and NSF were found by mass spectroscopy analysis to be present in the viral factories but not in the virion particle [58,59], whereas the SNARE protein was not found, consistent with its low expression profile. We therefore think that viral SNARE proteins, as they are not found in all giant viruses, do not act as viral membrane fusion proteins [20] but are probably used to interfere with the vesicle trafficking of the host cell. The expression profiles [57] suggest that all the factors of the core vesicle fusion machinery described above play roles during the viral reproduction cycle. We are aware that this is only conjecture at the moment and that in the future, the role of these trafficking factors in viral replication will need to be studied in more detail by examining their spatiotemporal patterns of expression using immuno fluorescence and confocal microscopy.

It should also be noted that we restricted our search to several core factors of the molecular machinery driving vesicle fusion in eukaryotic cells. Given the high number ofcore factors we found in the genomes of giant viruses, it is conceivable that other regulatory vesicle trafficking factors are present as well. For example, *Legionella pneumophila* expresses a serine protease that cleaves the SNARE protein syntaxin 17, blocking autophagy [60], whereas other factors interfere with the activity of different Rab proteins [61] In fact, in pathogenic intracellular bacteria, several factors regulating the activity of Rab proteins, e.g. GTPase activating proteins (GAPs) and guanine nucleotide exchange factors (GEFs), have been found. The search for such factors in the genomes of giant viruses should be pursued as well in the future to better understand how these viruses affect vesicular transport in the host cell.

## Methods

### Sequence search and classification

By using hmmscan from <u>HMMER</u> v3.2.1 [63], we searched for several membrane trafficking factors, namely SNARE [33], Rab [41], NSF[42], and SM proteins, with specific HMMs in viruses from the nr-database at NCBI as of 21 January 2020. We used a 10^-4^ expectation value cutoff. In an earlier study, we had already identified eukaroytic-like SNARE proteins in γ-proteobacteria of the order Legionellales and prototypic SNARE proteins in Asgard archaea using this approach [6]. To update our collection from γ-proteobacteria, we scanned the NCBI protein database again for bacterial sequences [6]. Next, we identified groups of similar sequence types by using the pairwise sequence similarity between all protein sequences as described earlier [6] using CLANS[64], which is based on the Basic Local Alignment Search Tool (BLAST)[65]. BLAST was also used to identify close eukaryotic homologues of viral proteins. Only e-values of ≤ 10^-15^ were used for clustering and visualization via a method from the Python package <u>networkX</u>. With this, we were able to define preliminary similarity groups of proteins. For a better understanding of their relationships, preliminary phylogenetic reconstructions using IQ-TREE [66] were carried out as well. To assess the presence of conserved domains and their arrangement within the similarity groups, we used SMART[67], and PFAM [68].

### Phylogenetic reconstruction

Phylogenetic reconstruction of the viral DNA polymerase B was carried out essentially as described[6]. Sequences were taken from [62]The alignment was built with Muscle with the standard parameters and gaps removed. The maximum likelihood tree was constructed with IQ-TREE [66] using the LG matrix[69] with a Γ-distribution for rate heterogeneity. We executed IQ-TREE with 1000 rapid bootstrap replicates[70]. Tree figures were made with FigTree v1.4.4 or iTOL [71]. The tree and alignments can be found at 10.5281/zenodo.5898721.

### Constructs, protein expression, and purification

All recombinant proteins were cloned into the pET28a vector, which contains an N-terminal, thrombin-cleavable His_6_-tag. The constructs for neuronal SNARE proteins from *Rattus norvegicus* have been described previously: the SNARE domain of syntaxin 1a (aa 180–262, Syx), a cysteine-free variant of SNAP-25b (aa 1–206, SN25), and synaptobrevin 2 (aa 1–96, Syb2) [27, 29]. Codon-optimized versions of the following SNARE protein sequences were synthesized and subcloned into the pET28a vector (GenScript): *Acanthamoeba polyphaga mimivirus*, YP_003987178.1, aa 1–204 (Mimi_Qbc), *Tupanvirus* deep ocean, QKU33625.1, aa 113-173 (TuVi_R), *Klosneuvirus* KNV1, ARF12037.1, aa 114–174 (KlVi_R, and ARF11934.1, aa 1-95 (KlVi_Qa). All proteins were expressed in the *Escherichia coli* strain BL21 (DE3) and purified by Ni^2+^-chromatography. After cleavage of the His_6_-tags by thrombin, the proteins were further purified by ion exchange chromatography on an Äkta system (GE Healthcare). The proteins were eluted with a linear gradient of NaCl in a standard buffer (20 mM Tris (pH 7.4) 1 mM EDTA) as previously described[6] [72,73]. The eluted proteins were 95% pure, as determined by gel electrophoresis. Protein concentrations were determined by absorption at 280 nm and the Bradford assay. SDS-PAGE was carried out as described by Laemmli. Non-denaturing gels were prepared and run in an identical manner to the SDS-polyacrylamide gels, except that SDS was omitted from all buffers [27, 29].

## Acknowledgements

This work was supported by the Swiss National Science Foundation (Grant 31003A_182732 to D.F. We thank the Division de Calcul et Soutien à la Recherche of the UNIL for access to the university’s computer infrastructure. We thank all members of the Fasshauer Laboratory for helpful discussions.

## Author contributions

E.N., D.K., and D.F. designed the study, performed the experiments, and analysed the data; D.F. wrote the paper; and all authors reviewed and commented on the manuscript.

## List of abbreviations

(NSF): N-ethylmaleimide-sensitive factor
(SNARE): soluble *N*-ethylmaleimide-sensitive factor attachment protein receptor
(SM): Sec1/ Munc18
(TMD): transmembrane domain
(HMM): hidden Markov model
(ER): endoplasmic reticulum
(GAP): GTPase activating proteins
(GEF): guanine nucleotide exchange factors

