## Supplemental Data for "Megaviruses contain various genes encoding for eukaryotic vesicle trafficking factors"

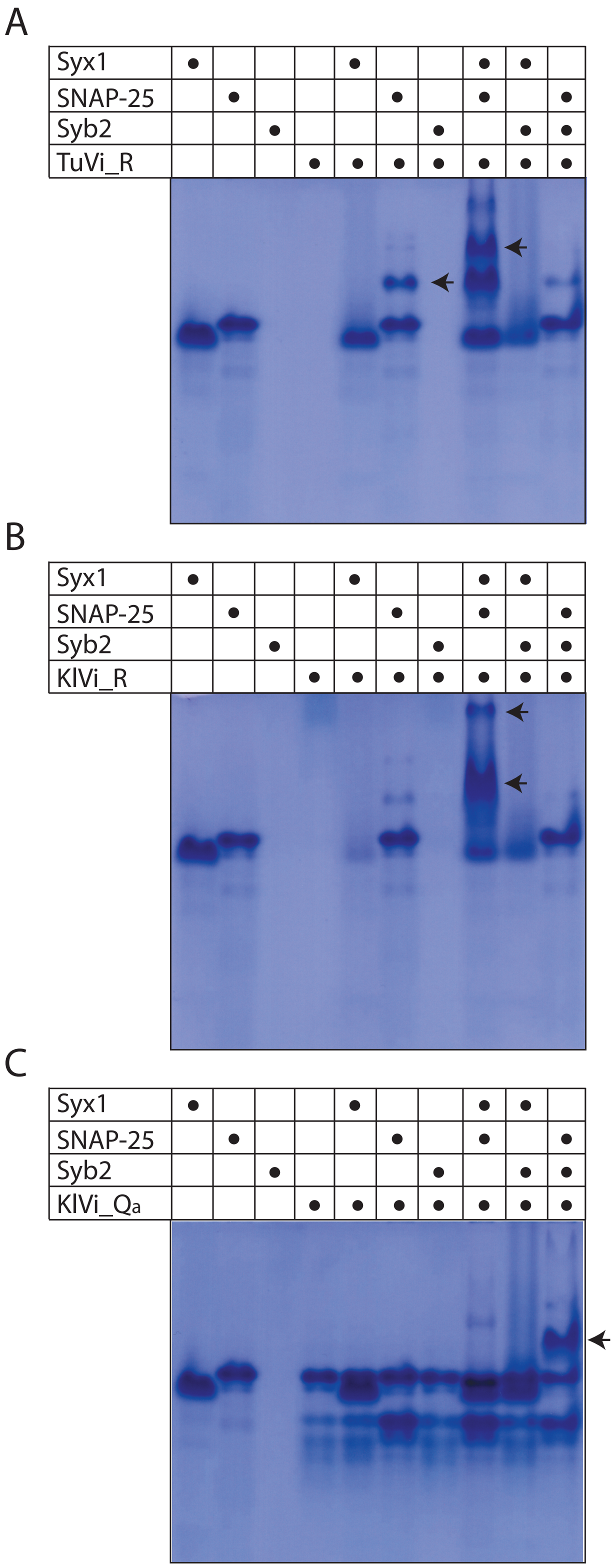

**Fig. S1. Formation of stable complexes between neuronal SNARE proteins and SNARE proteins from giant viruses monitored by non-denaturing gel electrophoresis.**
The proteins were incubated overnight at 4°C with equimolar ratios at ~15 µM concentration prior to non-denaturing gel electrophoresis, which separated native proteins and stable interactions by their charge only. Proteins were visualized by Coomassie Blue staining. An R-SNARE from *Tupanvirus* deep ocean (TuVi_R, QKU33625.1, A) formed a stable binary

complex with SNAP-25 alone and a stable ternary with SNAP-25 and Syntaxin 1 (Syx1). Complexes are indicated by arrows. An R-SNARE from *Klosneuvirus* KNV1 (KlVi_R, ARF12037.1, B) formed a stable with SNAP-25 and Syntaxin 1 and a Qa-SNARE from *Klosneuvirus* (KlVi_Qa, ARF11934.1, C) formed a stable complex with SNAP-25 and synaptobrevin 2 (Syb2).

**Table S1: List of bonafide SNAREs in γ-proteobacteria of the order Legionellales.** Four basic types, namely Qa-, Qb-, Qc-, and R-SNAREs, can be distinguished by their sequence profiles. Qbc-SNAREs consist of proteins with two different types of SNARE motifs, Qb- and Qc-motifs that are connected by a long linker. Sequences were clustered to construct groups for further inspection as described earlier [1].

| | Sequence ID | Sequence & SNARE motifs | Type | Species | Cluster ID |
|---|---|---|---|---|---|
| **R** | | | | | |
| | SNV31155.1 | MKSYALATQFNGESLQWHVPGNTYNPTTLFARKALGKWKKDIDDFVRQMQPGFCHCLYLDGYYIYGQKLGDKACVIVCDTELTIEQMRYLAYYLLNIGVDRETVAAHMEKYTRDEKVEQVKQELGEVKKIMIDNIDKVLERGERIEDLIKRTEGLADTSFLFHKKAKELNSCWPCTIF | R | *Legionella spiritensis* | 0 |
| | OEH47953.1 | MRCYALGTSFGDASFDWLISLSNSFLGGSFFEKQIKQTVEKYKEDINLYCSELKPNEVNWTKKGNIYIYAKRLVNDYCVVITDSPLTETQMYWLSFYLLKLQVDKDKIAPNIEKFTEDYKVVQVKKELEDVKQIMLENVEKVTLRGEKIDDLLEKTEQLEASSFQFKKNAEKLNSCWPSCVLL | R | *Legionella parisiensis* | 0 |
| | WP_058501541.1 | MKCYAIAVKFGAEKMEWVTAHSSLWTSVFFQNMEKYYELEVEKFLQGMKPGDQHIAQKNGYCLHAQRHMDDYCIIITDQLLSRGQLAHLTTYLLVLEEKVNKSIVFKDFEQYCSDRKLRQIKEELEETKKIMTDNIDKLLERGERIEQLIEKTEELSRNA | R | *Legionella israelensis* | 0 |
| | WP_058442445.1 | MKCFALGTRFGDAPFQWLTPGSNMTSFFAQKALTKHKQEIVAFCETLGTNETQGMQKDGYYFFIKRVGSDYCVVVTDTMLNEKQMNYLGWYLLRQSVSMSTVAADLEKHTKDFKVESIKQELAETQKIMLSNLDKVIERGERIEEVLAKSENLATSSFHFKQKAEELNSCWPSCVLI | R | *Legionella brunensis* | 0 |
| | KTD10104.1 | MRCYALGTCSGDGPFEWFISLSNSFFGSSFFEKQIKQAIEKHQAEITNYCADLNPNETYWMEKDGIYIYVRRVVDDYCGVIVDSKLDQKQITWLSLYLLKFKLEKKTIAANIEEFTQDYKVRKVQKEVDETMQIMLKNIEKMEKRGEALQTLVTKTQDLETSSFQFKKKAADLNSCWPSCTLI | R | *Legionella hackeliae* | 0 |
| | OGV52230.1 | MKCIALATHFGQKKTNWHIPSSNYSPSAFFTTRSLKKFEEDLRTQIIDLFEILQSGESIRFIEDMYFIHIQKLDNDCCAILCDTELTPQQMNYLSIALLSQQIPLKVIADNIEQYMQDYKTLALKADVEEVLNTMKHNMEKTLQRDIKIDDLVDKADKLEKHSIQFQRTVHEKTSSCWPSACNLL | R | *Legionellales bacterium RIFCSPHIGHO2_12_FULL_42_9* | 0 |
| | WP_058533815.1 | MKCFGLATKSGTNSFTWHIPGKGPFGFFGNLLPESIKKEITHFCDTQVHAGQIHYFNKNNYWFSVYKVNENCCIIAADCKLDSSQMSYLYLYLFDEEIPATTVASNLEKYTQNEKISDIKETLDETKKLMQNNIEQMLQRQEKIEDLAKRSEALAQGALSFKHKSEELNSCCILL | R | *Legionella saoudiensis* | 0 |
| | WP_058532341.1 | MKCFAIITRFEGEKATLHIPSNRIMLSLMASQIKKQLVKLEEIGEALEPGKYYFHTIGEIKIYGLKLGENYWAIACDSEITLEQQRNLTINLLFHKCSPQDVAQDLDQFIDKTHDPKIQKVQKELHETHAVLRETLEKMELRGEKLSDLVAKTEGLSQASFAFKEESERLNRCWPSCTLI | R | *Legionella rubrilucens* | 0 |
| | SNV46269.1 | MKCYAVGTRFGKTPFQWVIPGSDVTRFFAQRALETRKKEIISFCDELEDNETHRIQKDGYFVHIKKIGNDVCAIIVDKELSDKEMLYLSFHLLKANVEMSEIAENPDKYTEDPKIATIKRDLAETLQIMLENLEKTKLRGEKLEHLVATTQDLEASSFRFKKSAEDLNSCWPRFCTLI | R | *Legionella lansingensis* | 0 |

| | | | | | |
|---|---|---|---|---|---|
| | WP_081778127.1 (≈ SNV46269.1) | MIMKCYAVGTRFGKTPFQWVIPGSDVTRFFAQRALETRKKEIISFCDELEDNETHRIQKDGYFVHIKKIGNDVCAIIVDKELSDKEMLYLSFHLLKANVEMSEIAENPDKYTEDPKIATIKRDLAETLQIMLENLEKTKLRGEKLEHLVATTQDLEASSFRFKKSAEDLNSCWPRFCTLI | R | *Legionella lansingensis* | 0 |
| | WP_058526714.1 | MKCYAIITRFEGERATLHIPLNPIMLSLMGDQIKKHVAKLEEIGEALEPGKYFFDTIGEMRVYGLKLGGNYWAVACDSDMSLEQQRNLTLNLLFHKYPLPEVAQHLSQFMDKALDPKIQKFQQEIDETRDAALKALDSLDMRGEKISDLLQKTQSLSEASFAFKEKSEDLNRCWPCVIL | R | *Legionella erythra* | 0 |
| | ASQ47094.1 | MKCYALGTRFGDTSFDWLISVNSSFFGSGFFERQIKQSIEKHKDEIASYCSMMNPNETFGTKKEGFYLYITRLVHDYCVIVVDSELSEQQMRWLCFYLLKKKIEKNTVAANIEEFTQDYKITQIKSELHETTEIMKQNIEKLHLREEGLKKLVEKTEELETLTFHFNKKAKALNSCWPSCVLI | R | *Legionella clemsonensis* | 0 |
| | WP_094091873.1 (≈ ASQ47094.1) | MCQLLTWRIAMKCYALGTRFGDTSFDWLISVNSSFFGSGFFERQIKQSIEKHKDEIASYCSMMNPNETFGTKKEGFYLYITRLVHDYCVIVVDSELSEQQMRWLCFYLLKKKIEKNTVAANIEEFTQDYKITQIKSELHETTEIMKQNIEKLHLREEGLK | R | *Legionella clemsonensis* | 0 |
| | WP_108294219.1 | MKCYAIITRFEGEKATLHIPTNRIMLSLMATQIKKQVAKLEEIGEALEPGKYYSHTLGEMKIYGFKLGENYWAIACDSEITLEQQRTLTINLLFHKCALQDVAQDPDQFIDKAHDPKIKKVQKELDETHAVLLEAFQKMELRGEKLSELVDKTEGLSQLSFAFKEESERLNRCWPSCTLI | R | *Legionella taurinensis* | 0 |
| | SFL49537.1 | MRCYALGTRFGDTSFDWLISINSSFFGSGFFERQIKQSIEKHKDEIISYCSTMKPNETIGTKKEGFYIYLTRLVNDYCVVVVDNKLSEQQMRWLCFYLLKNKIEKNTVAANIEQFTQDYKVIKAQTAISETMQIMQENIGKLHTREEALQKLMKQTEELETLSFQFNKKAKELNSCWPSCVLI | R | *Legionella jamestownensis DSM 19215* | 0 |
| | WP_028387504.1 | MQNKDEQPISIATSSSSMASSSSTPARPPLICTGIARRFAVNPTESWIEHAGYGLFASPLGRAWVEQLEKKALELIDNYPYVISMNNYLVHFYRVGANWCAAALNRELSDGELRSLVHFLLYEKIPLATVAHTPKNYVTRIKATRDALDDTTEIMRDNLEKFLERREALERLLEDTEELKIETNRFYLSTTELNSCWPDKPWWLPSLSTISSMCQIL | R | *Legionella geestiana* | 0 |
| | WP_126339534.1 | MKCYALATQFNGESLQWHVPGNTYNPTTLFARKALGKWKKDIDDFVRQMQPGFCHCLYLDGYYIYGQKLGDKACVIVCDTELTIEQMRYLAYYLLNIGVDRETVAAHMEKYTRDEKVEQVKQELGEVKKIMIDNIDKVLERGERIEDLIKRTEGLADTSFLFHKKAKELNSCWPCTIF | R | *Legionella spiritensis* | 0 |
| | WP_143865370.1 | MKCYAIAVKFGAEKMEWVTAHSSLWTSVFFQNMEKYYELEVEKFLQGMKPGDQHIAQKNGYCLHAQRHMDDYCIIITDQLLSRGQLAHLTTYLLVLEEKVNKSIVFKDFEQYCCDRKLRQIKEELEETKKIMTDNIDKLLERGERIEQLIEKTEELSRNAKQFKHEAEELNRCWPSCTIL | R | *Legionella israelensis* | 0 |
| | WP_135061168.1 | MKCYAIAVKFGAEKMEWVTAHSSLWTSVFFQNMEKYYELEVEKFLQGMKPGDQHIAQKNGYCLHAQRHMGDYCIIITDQLLSRGQLAHLTTYLLVLEEKVNKSIVFKDFEQYCCDRKLRQIKEELEETKKIMTDNIDKLLERGERIEQLIEKTEELSISAKQFKHEAEELNRCWPSCTIL | R | *Legionella israelensis* | 0 |
| | WP_133137464.1 | MKCFGLATKSGNTPFTWHIQGNSLFGMFSKIFPESTKKEITDFCDTQIQPSQIHYFNKDSYWFSVYKVKENYCIVAADCKLDASQMSYLYLYLFDEKIPMVTAASNIEKYTQNEKINEIKETLEETKKLMHNNIELMLQRQEKIEDLAERSDALAKEAVSFKHKAQELNSCCTLL | R | *Legionella rowbothamii* | 0 |
| | WP_115301670.1 | MKCYALATCINSPKVTWSIPDSSTNSNGFFIRRFLGRIKEDISALLYHLPLDTCQGFQKEGYYLYGQRISSGFYGLACDTKLNEKQLTYLSFYLFSLHIEPDLIASDLSMHTQDQKIMHIKAELEKTKEIMKQNIEKILARGEQIEELVAHTEKLKEMSSQLSFKFHYKTKKEANDLPHFCNLI | R | *Legionella beliardensis* | 0 |
| | WP_133132450.1 | MKCYALAARFGEHPFQWVTTDNLLPTHNFFARQAINSYKQEIISFCEKLIPGEIQGTKKNGYFFYAARLGSDYCIAVLDTQLTETQMNFLSYYLLRLKLDLQTVANDLEKYTKDNKLEEINLQLAETRELMLENLEQILKRGESIERLLERSEELKNSSLSFKHESRKLTTCWPCNLI | R | *Legionella sp. W10-070* | 0 |

| | | | | | |
|---|---|---|---|---|---|
| | WP_131782847.1 | MKCYMLATGSHLSNITWITPNSSNQQGTFFTKKFFERIKLDI<br>LSFLENLKIDACYGFEREGYYLYGQKTSSGFYALACNVQLNK<br>RQLTYLCHHLFKCHLEPFLIAADLPKYTQDYQLLQAQAELEE<br>IKEIIRKNIEKVIVKGEKFESLMATVEEFKQRSSQFKHKLKM<br>KEPTSFEPLQNYPVTQAEVKLEETPEILIKKIEEMMIEIEPI<br>DNLLAKTEELDQISFKFKRKGKNNTNYHLQFCNLI | R | *Legionella gresilensis* | 0 |
| | WP_115330684.1 | MKCYMLATGSNLSNITWTTPNLSVNQSAFFTKNFFERIKLDI<br>LSFLENLKIDACYGCEREGYYLYGQKTSSNFYALACNIQLNK<br>KQLTYLYHHLFECHCKPALIAAELSKYVQDYQLLQAQAELEE<br>IQEIIRKNIERVIVKGEKFESLMAKVEELKQRSSQFKHKSKM<br>KELTSFEPQDYLETQAQVKVEETPEILLKKIEEMMIEIEPID<br>NLLAKTEELNQVSFKFKRKEKNNASSDLRFCNLI | R | *Legionella busanensis* | 0 |
| | OGT45220.1 | MRLYAIGIYEYTQEDSPQFQLKKKAPPNKGGFSGLFSTDPLE<br>NLEKHTLKALPPEQDVTFRATYKDEYHYVKRYPAEGIVIAIV<br>SRNKLEPLELAYLFSNIHTIYSKPNLAVTLDAIILNPLNYIA<br>RDIVTAQLKQKVQEIKEIMLENLEKLLDRGERIELLVRKTDY<br>LANGATSFKREAKKLNGCCHW | R | *Gammaproteobacteria bacterium RIFCSPHIGHO2_12_FULL_41_20* | 1 |
| | OGT53257.1 | MLYAIGIYTNGKRVAQAQKDKGLFSGNQLGSFEKHTLTSLPS<br>EMVENRAYYFPMKDEHHYVINYLQDNILIAIVSKTELEIKEL<br>TYLRTNIRTIYLRPNLNATLDDVIVNPLGYINRDILITRLKE<br>QAAEVKTIMLDNIEKVLQRGEQLEVLLDKTDRLRLDAITFKG<br>NAKKLNRCRC | R | *Gammaproteobacteria bacterium RIFCSPHIGHO2_12_FULL_42_13* | 1 |
| | OGT60900.1 | MLYAIGIYDYHDYMFIETDKPKPIQSAQTKNMLSLFNDNLLA<br>SLEKETLPLIHNSIIENSIIENDQTYYNSTNDEHHYFHVLTD<br>KNIIIAINSARKLDPYEIARLMINIEHIHMQPELARTNLQAV<br>INNPLGFTSCDIITKNMQVDIQDTKQIMQTNIQKLTERGDKL<br>AELEKTTFELSKNANKFSEGAKELNKCRC | R | *Gammaproteobacteria bacterium RIFCSPHIGHO2_12_FULL_43_28* | 1 |
| | WP_114833497.1 | MLTAIGIYQIDPYSKPFVFQKIMFAQADKSYFFQRNQLEKIE<br>KQSFLKLAGSLATDRFYCSQVDNEYHYIKMSADNIAIAVSSR<br>KELEKSEIAYLFANIRHIYMRNQNINQTLDNVIINPFGFTGK<br>DLLISHTQQNLDELKLELFDVLGKVLDRGNALEELQPKVIKL<br>NEASNKFRQAAERQRSCCRYW | R | *Aquicella lusitana* | 1 |
| | OGT42784.1 | MQIFCAVLYMANNQLLSASIDGSWWSFNAYEKNIRNHFPRVA<br>ELIKNQIGIHMLKWDRGFYYAFVFENKTAVLALDQTISDHAA<br>YSIYQHLQPITHVKELQTFISEPDEAIQSKADKIEAELKEVK<br>NIALNNINKILERGEKIESLVSQTSDLANQTVEFKKKSDHLN<br>RTCPGLFSVFSTISSYIWKTPAYEYHEIVHRRKNK | R | *Gammaproteobacteria bacterium RIFCSPHIGHO2_12_FULL_37_34* | -1 |
| | WP_114834964.1 | MKLFAIGLFHQNGTVEHIVTHEEYEALANAHESLASQIKTGI<br>YKNDVIYKIPAYGLYHYFKSSQGSIRILSSPENLDPTDPVLS<br>DQFSDINEIYLSTDHVVRYNELDKKIADPNYFISPVKRTRQG<br>VEETRQIMVNNIEKLLQRGAKLEELVEKTEELKQNTQSFRSK<br>AFKLNKNHTTVSEQAKAAAKKIFCCFFLPCCNGKPQGTAEER<br>ESFLRPNRSTIN | R | *Aquicella lusitana* | -1 |
| | KRG19470.1 | MTDPTIKPLTADDIKIQDMTETIDRTKRIMVDNIDQLLQRGE<br>KLEQLLDKTAQTEEKAQIFTQQARQIQIKARFENIAMTAAMI<br>GFILGGFYGLSAGIGLPMVAICGGIGGVVFYSVVWMFSGAIQ<br>SILKLPFFNLGFSPNIEKIEDESISVTKNFHPSLDYVKPNLI<br>YSNKHTVRCQLPAVSQAIDLEAQSMKKLTSRL | R | *Candidatus Berkiella cookevillensis* | 3 |
| | WP_057623293.1 (≈ KRG19470.1) | MPWIHSIICAVLIFSVKLKRCVMTDPTIKPLTADDIKIQDMT<br>ETIDRTKRIMVDNIDQLLQRGEKLEQLLDKTAQTEEKAQIFT<br>QQARQIQIKARFENIAMTAAMIGFILGGFYGLSAGIGLPMVA<br>ICGGIGGVVFYSVVWMFSGAIQSILKLPFFNLGFSPNIEKIE<br>DESISVTKNFHPSLDYVKPNLIYSNKHTVRCQLPAVSQAIDL<br>EAQSMKKLTSRL | R | *Candidatus Berkiella cookevillensis* | 3 |

| | | | | | |
|---|---|---|---|---|---|
| | OJV90465.1 | MKKNNQANERTKIKQIQQEIDETKKIMLQNIDKVIVRGRKIETLRDRSKEMQEKSQRFVKQAKHLKNKEKFYNLALEIVIVGAGVGAAYGVFAGYGWPLILTMAFLGGSAGYAVAWIWSGIQQKISSIFSYGKYIFSQGEVPTSFDASMNKPPLLKDKEALLSQYQNTANEVLSHYAAKEEPRLNNDKSKPRW | R | *Gammaproteobacteria bacterium 39-13* | 3 |
| **Qc** | | | | | |
| | SNV19632.1 | MKKNSSTKKILSSLSNTELQEQQELMMREQDEQLDILLKTVTRTHEIAEAIHKELTSQNKLIDGLNEDVEKTDGKVENTTKRVEALIPEVRSSCFLM | Qc | *Legionella pneumophila* | 2 |
| | WP_027219386.1 | MKKNSSTKKILSSLSNTELQEKQELMMREQDEQLDILLKTVTRTHEIAEAIQEELTSQNKLIDGLNEHVEKTDGKVKNTTKKVEALIPEVRSSCFLM | Qc | *Legionella pneumophila* | 2 |
| | SQG86500.1 | MKKNSSTKKILSSLSDTELQEKQELMMSEQDEQLDILLKTVTRTHEIAEAIQEELTSQNKLIDGLNEHVEKTDGKVKNTTKKVEALIPEVRSSCFLM | Qc | *Legionella pneumophila* | 2 |
| | WP_061637776.1 | MKKNSSTKKILSSLSNTELLEKQELMMSEQDEQLDILLKTVTRTHEIAEAIQEELTSQNKLIDGLNEHVEKTDGKVKNTTKKVEALIPEVRSSCFLM | Qc | *Legionella pneumophila* | 2 |
| | CZR11020.1 | MKKNSSTKKILSSLSDTELQEKQELMMREQDEQLDILLKTVTRTHEIAEAIQEELTSQNKLIDGLNEHVEKTDGKVKNTTKKVEALIPEVRSSCFLM | Qc | *Legionella pneumophila* | 2 |
| | PYB62981.1 | MKKNSSTKKILSSLSNTELLEKQELMMREQDEQLDILLKTVTRTHEIAEAIQEELTSQNKLIDGLNEHVEKTDGKVGNTTKRVETLIPEVRSSCFLM | Qc | *Legionella pneumophila* | 2 |
| | PQM70780.1 | MKKNSSTKKILSSLSNTELLEKQELMMSEQDEQLDILLKTVTRTHEIAEAIQEELTSQNKLIDGLNEHVEKTDGKVGNTTKRVETLIPEVRSSCFLM | Qc | *Legionella pneumophila* | 2 |
| | WP_061484634.1 | MKKNSSTKKILSSLSDTELQEKQELMMREQDEQLDILLKTVTRTHEIAEAIQEELTSQNKLIDGLNEHVEKTDGKVENTTKKVEALIPEVRSSCFLM | Qc | *Legionella pneumophila* | 2 |
| | ADG25747.1 | MKKNSSTKKILSSLSNTELQEKQELMMREQDEQLDILLKTVTRTYEIAEAIQEELTSQNKLIDGLNEHVEKTDGKVENTTKKVEALIPEVRSSCFLM | Qc | *Legionella pneumophila 2300/99 Alcoy* | 2 |
| **Qa** | | | | | |
| | WP_058512658.1 | MYKKIFKKDQPVSPSSTQQMLTFFSENEIDSGEQALRELEKRERTIREIERKIIDINQLFVDVAELVKDQGSLVNNIELSIEGAKEETAKGVIELEKAKKTQASTLCYLM | Qa | *Legionella santicrucis* | -1 |
| | WP_057625376.1 | MLGNGTPKLFSTSARKVSGDDKSSRKKKLTAETFLNAQHDDIEHVENGVTQLKSLSLDINELINNQNEMVEHIATNTEIGKENMNQGNENIEDAKRHHRHCCCTVL | Qa | *Candidatus Berkiella cookevillensis* | -1 |
| **Qbc** | | | | | |
| | OGT45367.1 | MRRALFEQGSQPDNKKLPESSNASTSHASIQQQLQQQREAKRAIAEEMLMSVANTNAQCAETLAELDRQGEQLARAHDKVRGVHEHLSMAEQHLHRMEHFFPSLSRPPQPPQPVADTASDRRKAQAPVVTSPANQPRQSSMEQSASSSDDAQLDMALEQITNGLQLLQNQSRVLGAELDRQNAKLGGLEQDVDTANSRLRQDNRRVRNLM | Qbc | *Gammaproteobacteria bacterium RIFCSPHIGHO2_12_FULL_41_20* | -1 |
| | WP_028380397.1 | MRMKKELFFQEKIQEINEVIEDSEASVERAARMARRTREIGADTVEALEGQSKKIQRIEKITENIQNTAEQAEESILNIKTYGLSRFFPPMPDFFNHPFRRKSKDEPPRHSTLTQSIKQELPRHHEPKLKTQYNEPVDRTLLDEKHLELLDKTNEGLDELGEIIDDLNQLAREQGKEITWQNEHLNKLNPKIEKATESINYETRRAGML | Qbc | *Legionella cherrii* | 4 |

| | | | | | |
|---|---|---|---|---|---|
| | SIR48573.1 | MRNKNELLFQKKIHEINEVIEDSAASVGRAVRMARNTREIGADTVEALEGQSKQIQKIEEITENIQNKAEQAEESILNIKTYGLSRFFPPIPDFLKHPFSRKSKDEPPRHSTLTQSIKQELPRHHEPKLKQQYNEPVDRTLLDEKHLELLGKTNEGLDELGEIIDDLNQLAREQGKEITWQNKHLDKLKPKIEMAKETVNYETGRAGML | Qbc | *Legionella (Fluoribacter) gormanii* | 4 |
| | WP_057622768.1 | MPKNKNFKKNRVLFDSAVIDTEEFDPETKEKIHTIVIDSNHSAHQILKIVNETKKIGEDTRKRLDQQGEQIGVIDRDVQELDYQAKKNKRTVKGINSVWWAIVHFFTPKCLKPQKPNFDDAVVEEDEKKNLAEISDFFTENTGTVTILFDERTKSVADDTSKVLSEANTAMHDLEFLSMGMNKKLRKQNKALDKIADKTDDINDNLRKTSKYARSYLGEPK | Qbc | *Candidatus Berkiella cookevillensis* | 5 |
| | KRG20005.1 (≈ WP_057622768.1) | MMPKNKNFKKNRVLFDSAVIDTEEFDPETKEKIHTIVIDSNHSAHQILKIVNETKKIGEDTRKRLDQQGEQIGVIDRDVQELDYQAKKNKRTVKGINSVWWAIVHFFTPKCLKPQKPNFDDAVVEEDEKKNLAEISDFFTENTGTVTILFDERTKSVADDTSKVLSEANTAMHDLEFLSMGMNKKLRKQNKALDKIADKTDDINDNLRKTSKYARSYLGEPK | Qbc | *Candidatus Berkiella cookevillensis* | 5 |

**Table S2: Overview of the distribution and types of vesicle trafficking proteins in giant viruses.**
Sequence IDs for viral vesicle trafficking factors are given. The table is related to Table S1, in which the presence and number of trafficking genes in viral genomes are illustrated.

| Clade | Virus | SNARE | | | | | Ras superfamily | | | | NSF | | SM protein |
|---|---|---|---|---|---|---|---|---|---|---|---|---|---|
| | | Qa | Qb | Qc | Qbc | R | Ras-& Rho-like | Ran-like | Rab-like | Rab5-like | Type1 | Type2 | |
| Mimiviridae & related viruses | *Samba virus* | | | | AHJ40280.1 | | | | | AHJ39970.2 | | AMK61911.1 | |
| | *Acantha. castellanii mamavirus* | | | | AEQ60861.1 | | | | | AEQ60397.1 | | AEQ60663.1 | |
| | *Acantham. polyphaga lentillevirus* | | | | | | | | | EJN40673.1 | | EJN40911.1 | |
| | *Hirudovirus strain Sangsue* | | | | AHA45183.1 | | | | | AHA45658.1 | | AHA45383.1 | |
| | *Acantham. polyphaga mimivirus* | | | | AKI79438.1, AKI80394.1 | | | | | AAV50487.1 | | YP_003986983.1 | |
| | *Mimivirus Bombay* | | | | AMZ03100.1 | | | | | AMZ02661.1 | | AMZ02920.1 | |
| | *Niemeyer virus* | | | | ALR84245.1 | | | | | ALR83777.1 | | ALR84066.1 | |
| | *Moumouvirus maliensis* | | | | | | | | | QGR53784.1 | | QGR53867.1 | |
| | *Moumouvirus australiensis* | | | | | | | | | AVL94639.1 | | AVL94725.1 | |
| | *Saudi moumouvirus* | | | | | | | | | AQN68133.1 | | AQN68231.1 | AQN68681.1 |
| | *Moumouvirus monve* | | | | | | | | | AEX63029.1 | | AEX62900.1 | AEX62303.1& AEX62302.1 |
| | *Acanthamoeba polyphaga moumouvirus* | | | | | | | | | YP_007354220.1 | | YP_007354307.1 | |
| | *Moumouvirus goulette* | | | | | | | | | AGF85510.1 | | AGF85417.1 | |
| | *Powai lake megavirus* | | | | | | | | | ANB50431.1 | | ANB50521.1 | ANB50942.1, ANB50531.1 |
| | *Mimivirus sp. SH* | | | | | | | | | AZL89530.1 | | AZL89444.1 | |
| | *Megavirus lba* | | | | | | | | | AGD92219.1 | | | AGD92819.1, AGD92328.1 |
| | *Megavirus chiliensis* | | | | | | | | | YP_004894365.1 | | YP_004894459.1 | YP_004894913.1,YP_004894469.1 |
| | *Bandra megavirus* | | | | | | | | | AUV58254.1 | | AUV58350.1 | AUV58800.1, AUV58362.1 |
| | *Megavirus courdo7* | | | | | | | | | AEX61416.1 | | AEX61533.1 | AEX61546.1 |
| | *Megavirus vitis* | | | | | | | | | AVL93647.1 | | AVL93737.1 | AVL94171.1, AVL93747.1 |
| | *Megavirus courdo11* | | | | | | | | | AFX92348.1 | | AFX92451.1 | AFX92968.1, AFX92464.1 |
| | *Mimivirus C* | | | | | | | | | AVG47149.1 | | AVG46139.1 | AVG46149.1, AVG46587.1 |
| | *Tupanvirus deep ocean* | QKU34194.1 | | | QKU33933.1 | QKU33625.1, QKU33594.1 | | | QKU34262.1, QKU35416.1 | QKU34328.1 | QKU34254.1 | QKU34179.1 | |
| | *Tupanvirus soda lake* | QKU35453.1 | | | QKU35182.1 | QKU34862.1, QKU34830.1 | | | QKU34156.1, QKU35523.1 | QKU35589.1 | QKU35516.1 | QKU35439.1 | |
| | *Satyrvirus sp.* | | | | AYV85399.1 | | | AYV85810.1 | AYV85451.1 | AYV85683.1 | AYV85780.1 | | |
| | *Klosneuvirus KNV1* | ARF11934.1 | | | | ARF12037.1 | ARF11765.1 | | ARF11168.1, ARF11569.1 | | ARF12416.1 | ARF12653.1 | |
| | *Fadolivirus* | | | | | QKF94280.1 | | | QKF93897.1 | | QKF94390 | | |
| | *Indivirus ILV1* | ARF09520.1 | | | | | | | ARF09856.1, ARF09964.1, ARF09892.1 | | | | |
| | *Barrevirus sp.* | AYV77044.1 | | | | | | | | | | | |
| | *Mimivirus LCMiAC01* | | | | | QBK88695.1 | | | QBK88830.1 | | | | |
| | *Mimivirus LCMiAC02* | | | | | QBK89254.1 | | | QBK89412.1 | | | | |
| | *Hokovirus HKV1* | | | | | ARF10343.1 | | ARF11123.1 | ARF10721.1 | | ARF10712.1 | | |
| | *Bodo saltans virus* | ATZ80326.1 | ATZ80812.1 | | | ATZ81026.1 | | | ATZ80205.1, ATZ80809.1, ATZ80829.1, ATZ81046.1, ATZ81047.1 | | ATZ80361.1 | | |
| | *Terrestrivirus sp.* | AYV76306.1 | AYV76603.1, AYV75536.1 | AYV75598.1 | | AYV76397.1 | | | AYV75142.1, AYV75191.1, AYV75487.1, AYV75604.1, AYV75810.1, AYV75892.1, AYV76027.1, AYV76231.1, AYV76278.1, AYV76633.1, AYV76634.1 | | AYV75575.1 | | AYV76567.1, AYV75872.1 |
| | *Harvfovirus sp.* | AYV80658.1 | | | | AYV80658.1 | | | AYV80695.1, AYV80804.1, AYV81889.1, AYV81890.1, AYV81891.1 AYV81892.1, | | | | |
| | *Hyperionvirus sp.* | | | | | AYV84002.1 | AYV84145.1 | | AYV83648.1, AYV83649.1, AYV83650.1, AYV83657.1, AYV84174.1, AYV84774.1 | | AYV83750.1 | | AYV82503.1 |
| | *Catovirus CTV1* | ARF09111.1 | ARF08857.1 | ARF09317.1 | | ARF08579.1 | | | ARF08354.1, ARF08666.1, ARF08702.1, ARF09118.1 | | | | ARF08705.1 |
| | *Edafosvirus sp.* | | AYV78627.1 | AYV78098.1, AYV78103.1 | | AYV77932.1 | | | AYV77911.1, AYV77912.1, AYV77913.1, AYV77914.1, AYV78041.1, AYV78105.1, AYV78414.1, AYV78944.1 | | AYV78062.1 | AYV78487.1 | |
| | *Yasminevirus sp. GU-2018* | VBB19031.1, VBB17642.1, VBB18316.1 | VBB19035.1 ,VBB17939.1 | VBB19034.1, VBB19046.1 | | VBB18571.1 | | | VBB17603.1, VBB18222.1, VBB18224.1, VBB18244.1, VBB18303.1, VBB18902.1, VBB19039.1 | | VBB17791.1 ,VBB18968.1 | VBB18892.1 | VBB19033.1 |
| | *Faunusvirus sp.* | AYV79275.1 | | AYV79625.1 | | AYV79271.1 | | | AYV79135.1 | | | | |
| | *Mimiviridae sp. ChoanoV1* | QDY52074.1 | | QDY51946.1 | | | | | QDY51751.1, QDY52179.1 | | | | |
| | *Chrysochromulina ericina virus* | YP_009173670.1 | | | | | | | | | | | |
| | *Homavirus sp.* | | | | AYV82099.1 | | | | AYV82031.1, AYV82032.1, AYV82395.1 | | | | |

| | | | | | | | | | | | |
|---|---|---|---|---|---|---|---|---|---|---|---|
| | *Megaviridae environmental sample* | | | QFG73916.1 | | | | | QFG73830.1, QFG74315.1, QFG74977.1 | | |
| Unclassified | *Sylvanvirus sp.* | AYV87155.1, AYV87038.1 | AYV87113.1 | AYV87104.1 | | AYV87231.1 | | | | | AYV86903.1 |
| | *Acanthamoeba castellanii medusavirus* | | | | | | | BBI30496.1 | | | |
| Pythoviridae | *Pithovirus LCDPAC01* | | | | | | | QBK84778.1 | QBK84575.1 | | |
| | *Pithovirus LCDPAC02* | | | | | | | QBK84941.1 | | | |
| | *Pithovirus LCPAC001* | | | | | | | QBK89590.1, QBK89605.1, QBK89606.1, QBK89566.1 | | | |
| | *Pithovirus LCPAC101* | | | | | | | QBK90090.1 | | | |
| | *Pithovirus sibericum* | | | | | | YP_009001039.1, YP_009001038.1 | YP_009001029.1, YP_009001018.1 | | | |
| | *Pithovirus LCPAC103* | | | | | | | QBK90332.1, QBK90331.1, QBK90333.1 | | | |
| | *Pithovirus LCPAC201* | | | | | | | QBK90994.1 | | | |
| | *Pithovirus LCPAC202* | | | | | | | QBK91324.1 | | | |
| | *Pithovirus LCPAC302* | | | | | QBK91448.1 | | | QBK91453.1 | | |
| | *Pithovirus LCPAC304* | | | | | | | QBK92074.1 | QBK91707.1, QBK92208.1, QBK91734.1 | | |
| | *Pithovirus LCPAC401* | | | | | | | QBK92446.1 | | | |
| | *Pithovirus LCPAC403* | | | | | | | QBK93151.1 | | | |
| | *Pithovirus LCPAC406* | | | | | | | QBK93913.1 | | | |
| | *Orpheovirus IHUMI-LCC2* | | | | | YP_009448950.1 | YP_009449015.1 | | YP_009448845.1 | | |
| Iridoviridae | *Iridovirus LCIVAC01* | | | | | | | QBK85217.1, QBK85301.1 | | | |
| | *Scale drop disease virus* | | | | | | | | | | YP_009163787 |
| | *Lymphocystis disease virus Sa* | | | | | | | | | | YP_009342207.1 |
| | *Lymphocystis disease virus - isolate China* | | | | | | | | | | YP_073712.1 |
| | *Lymphocystis disease virus 2* | | | | | | | | | | BCB67535.1 |
| | *Lymphocystis disease virus 4* | | | | | | | | | | QHR78442.1 |
| Marseilleviridae | *Marseillevirus LCMAC102* | | | | | | QBK86376.1,QBK86444.1 | QBK86593.1 | | | |
| | *Marseillevirus LCMAC101* | | | | | | QBK85941.1 | | QBK85506.1 | | |
| | *Marseillevirus LCMAC202* | QBK88261.1 | | | | | | | QBK88238.1 | | |
| | *Marseillevirus LCMAC201* | QBK87296.1 | | | | | | QBK87254.1 | QBK87510.1 | | |
| Phycodnaviridae | *Emiliania huxleyi virus 164* | | | | | AHA55712.1 | | | | | |
| | *Emiliania huxleyi virus 18* | | | | | AHA54185.1 | | | | | |
| | *Emiliania huxleyi virus 99B1* | | | | | CAZ69436.1 | | | | | |
| | *Emiliania huxleyi virus 88* | | | | | AEP15042.1 | | | | | |
| | *Emiliania huxleyi virus 84* | | | | | AEO97660.1 | | | | | |
| | *Emiliania huxleyi virus 145* | | | | | AHA54679.1 | | | | | |
| | *Emiliania huxleyi virus 156* | | | | | AHA55231.1 | | | | | |
| | *Emiliania huxleyi virus 86* | | | | | YP_293857.1 | | | | | |
| | *Emiliania huxleyi virus 203* | | | | | AEO98116.1 | | | | | |
| | *Emiliania huxleyi virus 208* | | | | | AEP16122.1 | | | | | |
| | *Emiliania huxleyi virus 207* | | | | | AEP15645.1 | | | | | |
| | *Emiliania huxleyi virus 201* | | | | | AET97989.1 | | | | | |
| | *Emiliania huxleyi virus 202* | | | | | AET42546.1 | | | | | |
| Retroviridae | *Kirsten murine sarcoma virus* | | | | | | CAA80675.1, P01117 | | | | |
| | *LNras*SN acutely transforming retrovirus* | | | | | | ADI24415.1 | | | | |
| | *Moloney murine sarcoma virus* | | | | | | AAA46575.1, P23175.1, P01113.1 | | | | |
| | *Havey murine sarcoma virus* | | | | | | YP_009507789.1, P01115 | | | | |
| | *Rat sarcoma virus* | | | | | | P01114 | | | | |
